# A Bayesian nonparametric semi-supervised model for integration of multiple single-cell experiments

**DOI:** 10.1101/2020.01.14.906313

**Authors:** Archit Verma, Barbara Engelhardt

## Abstract

Joint analysis of multiple single cell RNA-sequencing (scRNA-seq) data is confounded by technical batch effects across experiments, biological or environmental variability across cells, and different capture processes across sequencing platforms. Manifold alignment is a principled, effective tool for integrating multiple data sets and controlling for confounding factors. We demonstrate that the semi-supervised t-distributed Gaussian process latent variable model (sstGPLVM), which projects the data onto a mixture of fixed and latent dimensions, can learn a unified low-dimensional embedding for multiple single cell experiments with minimal assumptions. We show the efficacy of the model as compared with state-of-the-art methods for single cell data integration on simulated data, pancreas cells from four sequencing technologies, induced pluripotent stem cells from male and female donors, and mouse brain cells from both spatial seqFISH_+_ and traditional scRNA-seq.

Code and data is available at https://github.com/architverma1/sc-manifold-alignment

## Introduction

A variety of single cell technologies allow biologists to measure gene expression in individual cells. Recent advances have reduced costs while improving throughput, leading to data consisting of thousands or even millions of cells and tens to tens of thousands of genes (1). This level of granularity opens up new insights not available from bulk RNA sequencing: the discovery and characterization of cell populations, the changing profiles of gene expression across development, and the cellular response to stimuli among others. Projects such as the Human Cell Atlas, the Human Tumor Atlas, and Tabula Muris aim to capture the entire space of cell types and states across tissues and conditions (2–4).

As the complexity and size of single cell projects have increased, researchers need to sequence multiple batches (5), consortia need to compile data from various member labs (2), data have to be combined across old and new technologies (6), and sample heterogeneity continues to grow (7). New methods are constantly developed to improve sequencing or add additional information about cells, such as spatial patterning (1). The increase in sample size improves the power of analyses (8, 9); however, the integration of multiple single cell data sets is often confounded by batch effects and other conditions that differ across cells. Several factors can lead to differential expression patterns across experiments: i) the use of different sequencing technologies and protocols, ii) varying environmental conditions and preparation methods across labs, and iii) different cellular characteristics including cellular environment (5, 7). Even technical replicates in scRNA-seq exhibit substantial variability (7).

The development of efficient, principled computational methods for correcting batch effects and integrating multiple experiments is thus critical to enable downstream analysis of single cell data sets. A simple solution has been to reuse tools developed for bulk RNA sequencing. Limma’s *removeBatchEffect* fits a linear model to regress out batch effects in bulk RNA-seq data, but can be used for scRNA-seq as well (10). ComBat adds empirical Bayes shrinkage of the blocking coefficient estimates to correct for batch (11). These and other bulk methods generally assume linear effects and fixed populations, conditions that are unlikely to hold when analyzing scRNA-seq data. In response, several methods have been developed to correct for batch effects in scRNA-seq. One method uses differences in pairs of mutual nearest neighbors (MNN) between cosine-normalized expression levels across experiments to calculate batch effect vectors – or the convex combination of each pair of mutual nearest neighboring cells – that can be used to project the cells onto a shared latent space (9). A previous release of Seurat, a single cell analysis R package, used a modified canonical correlation analysis (CCA) framework to remove batch effects (6).

A complete data integration procedure should provide: 1) a mapping between high and low dimensional spaces that removes unwanted variation; 2) uncertainty estimates in the alignment of possibly nonlinear manifolds; 3) reference-free regularization that preserves variation from sources other than batch; and 4) robust alignment when portions of the sub-spaces are not shared. Neither MNN nor CCA are probabilistic, and both struggle when populations are not shared across experiments. In addition, MNN and CCA can only correct for single categorical variables; in contrast, linear approaches allow complex covariates to be corrected (10, 11). Bulk methods, on the other hand, fail to account for nonlinear patterns in genetic expression and variance in the cell populations across samples.

Here we propose and demonstrate the use of robust semi-supervised Gaussian process latent variable models (12–14) to estimate a manifold that eliminates variance from unwanted covariates and enables the imputation of missing covariates for many types of metadata. We use a robust t-distributed Gaussian process latent variable model (tGPLVM) to account for the overdispersed and noisy observations in single cell data. Then, we allow the introduction of fixed covariates that encode known meta-data to estimate a shared low-dimensional nonlinear manifold; we refer to this semi-supervised tGPLVM as sstGPLVM. The fitted manifolds can then be collapsed across the fixed covariates to remove the effects of those covariates, and the projections can be mapped back to a high-dimensional expression matrix for downstream analysis. Similarly, missing meta-data or expression counts can be imputed from the estimated manifolds. We demonstrate this model’s applicability to integrating data across modalities and biological conditions including batch correction, sex differences, and spatial transcriptomics.

## Methods

### Description of the sstGPLVM

The Gaussian process latent variable model (13, 15) posits that each feature of high-dimensional observations, *Y* ∈ ℛ^*N×P*^, is generated from a Gaussian process (GP) projection of a lower dimensional representation of the samples *X* ∈ ℛ^*N×Q*^ plus some statistical noise. Traditionally, the Gaussian process has been assumed to have a Gaussian kernel, imposing a smooth manifold, and to have normally distributed noise. Previous work on gene expression data has challenged both of these standard assumptions (12, 16). In particular, we have seen that the manifold is not particularly smooth, leading to the use of a Matérn kernel in the Gaussian process. Similarly, the noise model is not well captured by the Gaussian distribution, but the heavy-tailed Student’s t-distributed error improves the performance of GPs on expression data, leading to the tGPLVM (12).

We expand on ideas behind the tGPLVM by adding *K* fixed variables that capture known covariate information about the cells possibly contained in the meta-data, such as batch, spatial location, or donor sex. This information can be encoded by a one-hot vector, such as batch, or a continuous variable, such as (*x, y*) coordinates representing spatial location, leading to *X* ∈ ℛ^*N×*(*Q*+*K*)^ low dimensional space. In this model, covariates can also include missing data if covariate information is only available for a limited number of cells. With these estimated low-dimensional embeddings, we can then both identify the differences in cells and gene expression across the fixed covariates and also control for the effects of the fixed covariates by projecting the manifolds along the fixed dimensions, e.g. drawing counts for all cells as if they came from the same experiment.

### sstGPLVM Generative Model

Given *N* observations across multiple data sets *Y*_1_, *Y*_2_, … with *P* shared features, we assume that the data come from a joint low-dimensional latent space 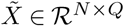 with a Gaussian prior:

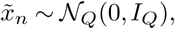

where *n* represents the sample index (*n* = 1 : *N*) and *Q* is the number of latent dimensions. For each data point there also exists *K* fixed variables *z*_*n*_ ∈ ℛ^*K*^. These fixed variables may represent a location in space, the batch from which the sample is derived, or any other known information or meta-data about the sample. We use binary, one-hot encodings to represent sample, batch, or individual in this paper. The fixed variables may also be mixes of known and missing values in the case that the relevant information is available for only some of the data types (e.g., (*x, y*) coordinates are available for seqFISH_+_ data but not for scRNA-seq data). We concatenate the latent variables and fixed variables: 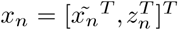, and the total dimension of the low-dimensional projection is then *Q* + *K*. The noiseless high-dimensional observations of each feature across samples, *p* = 1 : *P*, have a Gaussian process prior parameterized by the fixed and latent variables as previously described (12):

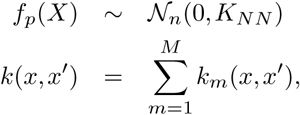

where *K*_*NN*_ represents the *N* × *N* covariance (Gram) matrix defined by *k*(*x, x*′) for *M* different basis kernels. We model the residuals with a heavy-tailed Student’s t-distribution:

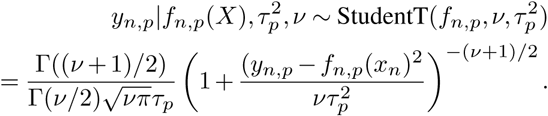

Throughout, we set the degrees of freedom *ν* = 4 based on prior work (12). Each feature-specific scale of the Student’s t-Distribution, 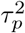, is a hyperparameter optimized during inference. We use a flexible sum of Matérn 1/2 and Gaussian kernels (*M* = 2) to allow for heterogeneous manifold topology, and automatic relevance determination (13) to estimate the importance of each dimension:

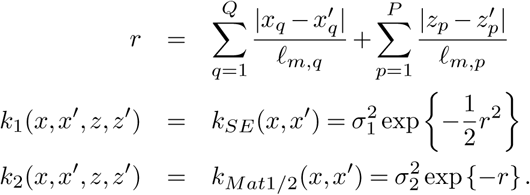

### Inference

We fit the model with black box variation inference (BBVI) (17) with the same variational distributions as used with fully unsupervised tGPLVM (12). BBVI was implemented in Edward (18) and Tensorflow (19). Iterations were run using the log_2_(1 + *Y*) transformation of the entire cell by gene count matrix unless otherwise specified on a Microsoft Azure 16 vCPU 224 GB H16m high performance computing cloud machine.

### Simulated data

We simulated two-dimensional data with 8 clusters representing cell types across two batches of sizes *N*_1_ = 300 and *N*_2_ = 200. Not all clusters were in both batches. The low-dimensional data were linearly projected up to 250 dimensions by a random matrix drawn from a Gaussian with variance *σ* = 5. Batch “effects,” nonlinear transformations, were added to each batch: batch one had a quadratic function of the cell’s latent position added to the high dimensional counts, and batch two had a cubic function of the cell’s latent position added to the high dimensional counts. The feature (“gene”)-specific coefficients were drawn from a Gaussian distribution with variance ranging from 1 to 5, increasing the average batch effect magnitude. Finally, we added Gaussian noise to each observation with mean zero and variance one.

The model was fit for each simulation for 1000 iterations. We then computed a distance between the estimated and simulated latent space. First, the pairwise distance matrix of the cell embeddings in both the estimated and simulated low-dimensional space is calculated. Each row is normalized to sum to one. We then calculate the 2-Wasserstein distance for each row, or cell, with the corresponding normalized distances in the true latent space, and we compute the average across all cells. This metric penalizes points that are distant in the true embedding for being close in the estimated embedding.

To compare to Seurat’s CCA (6), we estimated a joint representation of both batches using all features, no normalization, and two canonical components. To compare to MNN, we performed fastMNN in scater (9) with all features and calculated the distance matrix of the high-dimensional “corrected” observations that the function returns. To compare to linear methods, we first fit a linear regression model between a binary batch indicator variable and the observations. We then took the first two principal components of the residuals for use in calculating the 2-Wasserstein distance. To compare to sequential estimation of the known and latent effects, we also fit a Gaussian process regression between a binary batch indicator variable and the observations, followed by fitting a two-dimensional GPLVM with GPy. To determine the magnitude of the fixed effects, we calculated the 2-Wasserstein distance metric between the noisy observations and the true low-dimensional space as a benchmark for performance.

### Correcting batch effects across pancreas cells

We fit sstGPLVM to data from four pancreas data sets: GSE81076 (CEL-seq) (20), GSE85241 (CEL-seq2) (21) and GSE86473 (SMART-seq2) (15) and from ArrayExpress accession number E-MTAB-5061 (SMART-seq2) (22). We use the data as processed by scater in R following the pipeline from a previous study of the data (9) before correction. We fit our model for 500 iterations.

Based on hyperparameter optimization for the number of dimensions (Additional File 1), we used the five most important dimensions as determined by kernel length scales for downstream analysis. We performed ten repeats of K-means clustering with between five and ten clusters to compare to ground truth cell type labels provided using normalized mutual information (NMI) and adjusted rand score (ARS) from scikit-learn (23). This analysis was also performed to an expression matrix produced by MNN and a PCA embedding with ten dimensions after removing batch effects with ordinary least squares linear regression. We also used calculated the variational posterior (*q*_*f*_) (12) to estimate the high dimensional counts as if all the cells had been sequenced in the first batch, on which we performed Walktrap clustering in the scater R package (9). To understand the relationship between cell type heterogeneity and uncertainty in the estimated embeddings, we performed linear regression between the average uncertainty, quantified by the variance of the variational posterior of the embedding (*q*_*x*_), estimated across the five latent dimensions and the number of nearest neighbors out of twenty nearest neighbors that were of the same cell type.

### Learning sex specific manifolds from induced pluripotent stem cells (iPSCs)

We fit sstGPLVM to scRNA-seq data of iPSCs from 53 Yoruban individuals (8, 24). We fit a model with ten latent dimensions along with sex information for 500 iterations, as well as a model with only latent dimensions and no sex information. We fit a logistic regression model with scikit-learn to predict sex using the latent embedding to quantify how separable the manifold is with and without the inclusion of sex information in the latent space. We use the noiseless counts in the posterior variational distribution (*q*_*f*_) (12) to impute the counts of cells as if they were the opposite sex. We calculate the mean of the variational noiseless counts using the same fixed variable for all cells to indicate which sex to impute as (0 for female, 1 for male).

### Aligning scRNA-seq with seqFISH_+_ spatial mappings

We fit sstGPLVM to expression data from seqFISH_+_ data (25) without log normalization and the log count matrix from a scRNA-seq experiment both for mouse brain samples (26). The sstGPLVM has two latent dimensions as well as three fixed dimensions: the (*x, y*) coordinates of the cells, and the batch. However, the (*x, y*) coordinates are only available for the cells from seqFISH_+_. For the remaining cells from scRNA-seq, the (*x, y*) coordinates in low-dimensional space are set as variables to be learned during inference. We fit sstGPLVM with data from the olfactory bulb for 1000 iterations; we observed that after one hundred iterations the fit was stable and therefore fit the model to data from the cortex and sub-ventricular zone for only one hundred iterations. For cellular spatial analysis, we first filtered out cells that were not located inside the bounds of the seqFISH_+_ data. We then found the 15 nearest neighbors of each cell and identified the mutual nearest neighbors in order to quantify enrichment for neighboring cell types. For visualization, we then normalized the cell type proportions of the neighbors to sum to one and divided by the proportion of cell types across all cells embedded inside the seqFISH_+_ (*x, y*) coordinate space to identify enrichment beyond what is expected under the null distribution of uniform placements.

## Results

### Comparison of sstGPLVM versus related methods on simulated data

First, we simulated data with batch effects to verify the ability of different methods to recover unified latent spaces across multiple data modalities. We started with a two dimensional latent space that included samples from eight clusters in this low-dimensional space split across two batches with different cluster proportions (see Methods for full description). Some clusters were only present in a single batch. We linearly projected the data into a 250 dimensional feature space, and we added nonlinear batch effects to both batches: a parabolic function of the position in the latent space to batch one and a hyperbolic function of the position in the latent space to batch two. Finally, we added Gaussian noise. The magnitude of the gene-specific batch effects was drawn from a normal distribution with zero mean but increasing variance from one to five.

We fit our model to the noisy observations with batch encoded as a fixed variable. For comparison, we performed correction with Seurat (CCA) (6) and Mutual Nearest Neighbors (MNN) (9). We also created three benchmark comparisons: the uncorrected noisy observations (Batch), a PCA projection after using linear regression to correct batch effects (PCA), and a GPLVM embedding after using Gaussian process regression to correct batch effects (GPy). To evaluate the fit of each approach, we use a scale-invariant comparison of the simulated latent space to the estimated latent space based on the Wasserstein-2 distance (see Methods for details), with a smaller distance indicating closer manifolds and a more accurate embedding to the simulated truth.

Intuitively, we find that, as the magnitude of the batch effects increases, the distance between the true latent space and the observations increases (Figure 1). Linear correction with PCA slightly improves the fit, but follows the trends of the uncorrected observations closely. sstGPLVM has the lowest distance to the true space of all the methods including the benchmark approaches across batch effect magnitudes. Sequential correction first with Gaussian process regression and then with a nonlinear latent variable model performs worse than joint learning with sstGPLVM, suggesting that estimation of the manifold with fixed covariates allows for better allocation of variance across the unknown and fixed dimensions. MNN and CCA perform similarly to each other, and the distances of the fit are surprisingly consistent across magnitudes of batch effects. In particular, they tend to over-correct at the lower magnitudes, as indicated by their distance from simulated truth being larger than the uncorrected observations. The superior performance of sstGPLVM at all magnitudes, while remaining agnostic about the nature of the batch effects on the transcription data, suggests that joint semi-supervised learning most accurately integrates multiple data sets with known batch.

**Fig. 1.**
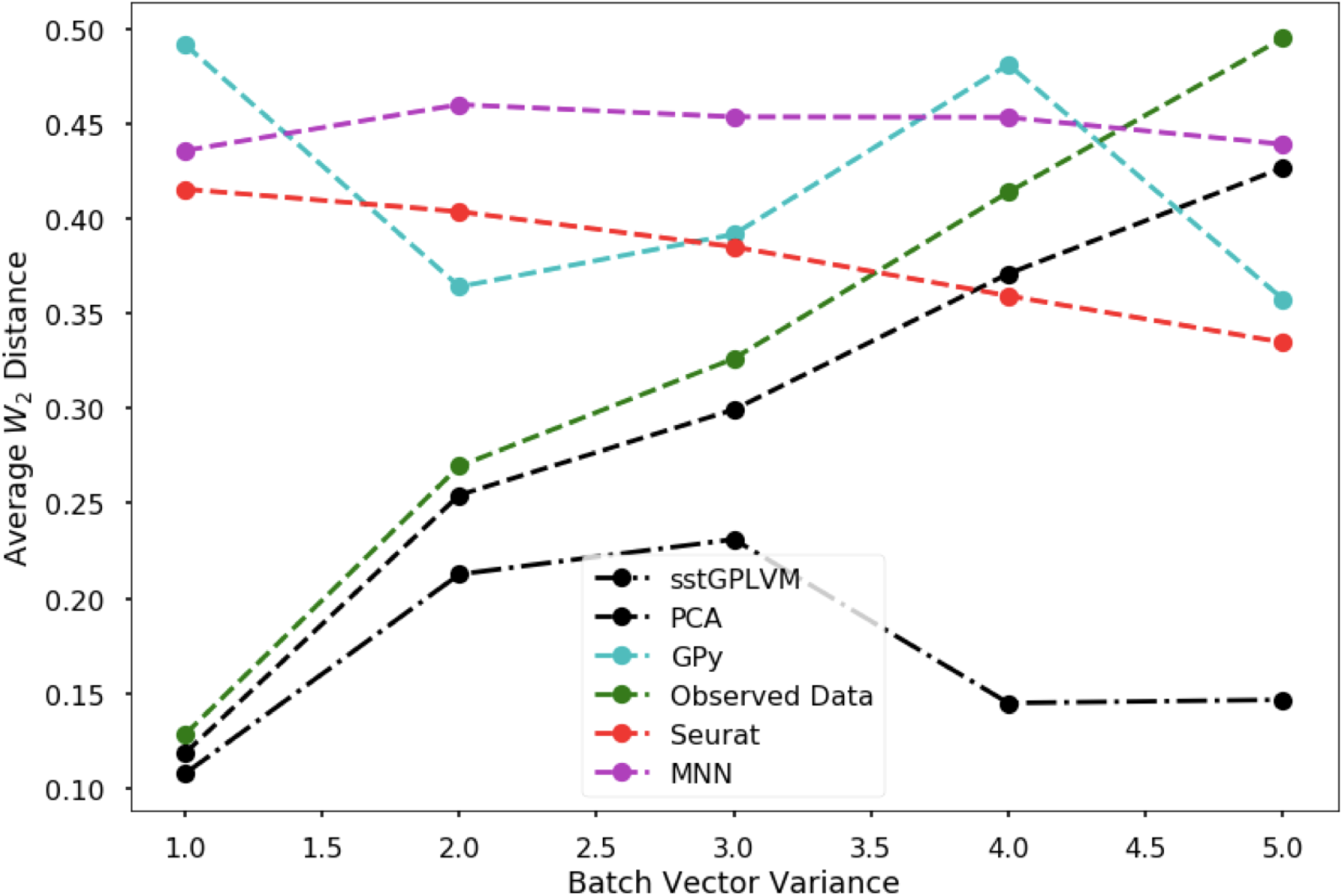
Performance on simulated data. Average 2-Wasserstein distance of estimated latent space to simulated low-dimensional space for multiple methods.

### Correcting batch effects in pancreas cells

Next, we tested the ability of our model to correct batch effects in scRNA-seq data from different platforms. We fit sstGPLVM to the four pancreas data sets analyzed in the MNN paper (15, 20–22). We evaluated the fit by performing ten repeats of K-means clustering of the projected cells with five to ten clusters and comparing to existing cell type labels with normalized mutual information (NMI) and adjusted rand score (ARS). Our sstGPLVM performs comparably to the original MNN analysis and outperforms simple linear correction (Table 1).

**Table 1.** Clustering scores for pancreas data. The average normalized mutual information (NMI) and adjusted rand score (ARS) over ten k-means clustering repeats with 5 ≤ *K* ≤ 10 and the ARS for Walktrap clustering.

|  | Latent Space, K-Means, NMI | Latent Space, K-Means, ARS | Gene Space, Walktrap, ARS |
| --- | --- | --- | --- |
| sstGPLVM | 0.42 | 0.34 | 0.44 |
| MNN | 0.45 | 0.42 | 0.50 |
| Linear Regression + PCA | 0.23 | 0.13 | NA |

Next, we projected the data back into the high dimensional (gene) space, assuming the data came from the same batch, as might be done to perform gene-level imputation. We tested the imputed expression levels with the scater pipeline’s Walk-trap clustering (9). Though the clustering scores are slightly lower (Table 1), the sstGPLVM allows us to visualize which cells are confidently embedded and which are more uncertain (Figure 2). Observing the latent space, we can see that cells that are in more mixed areas of the manifold are also more uncertain in their mapping. Linear regression between the number of nearest neighbors that are the same cell type and the uncertain in the cell embedding revealed a clear negative correlation (*R* = −0.29, *p* ≤ 2.2 × 10^−16^). The addition of this uncertainty information can be used to inform decision making in downstream analysis and future experimentation. Many analytical methods are used post-batch correction without interrogation of the quality of the batch correction. The uncertainty estimates from sstGPLVM provide a measure of quality control that can prevent the propagation of errors downstream.

**Fig. 2.**
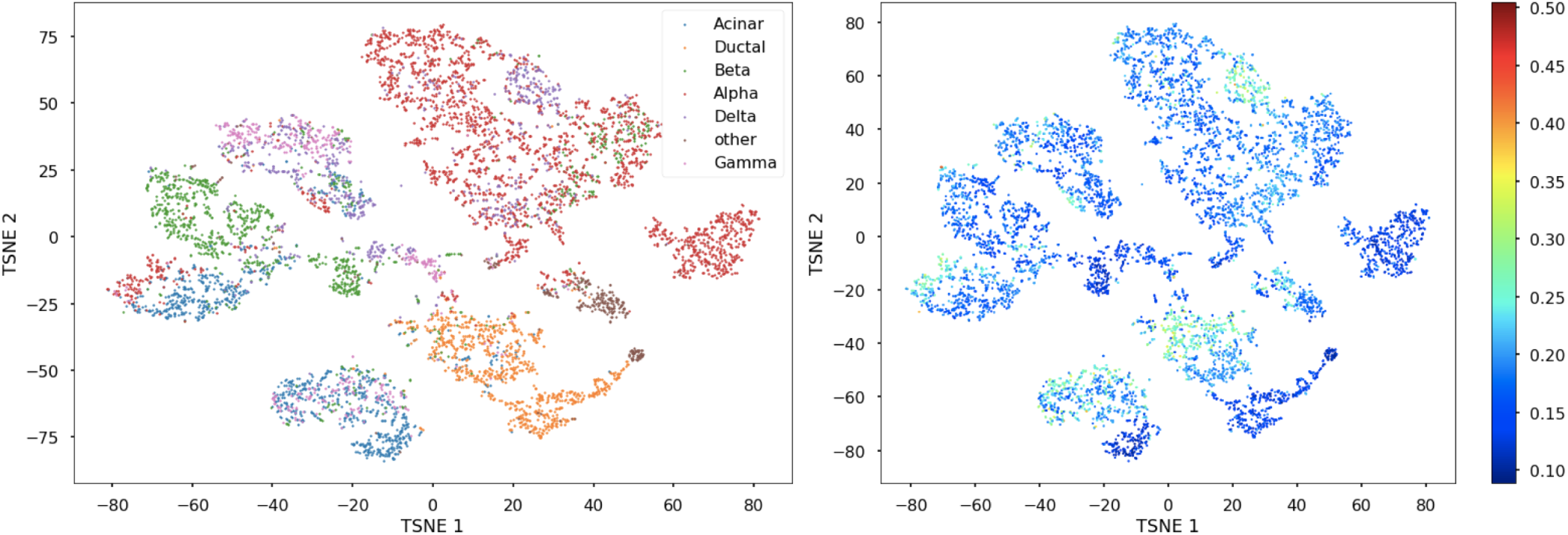
Pancreas cells integrated from four sources. T-SNE embedding of learned latent space labeled by (left) cell type, and (right) uncertainty of the embedding.

### Exploring sex manifolds in iPSCs

Next, we show that sstGPLVM is able to correct for biological covariates that confound results. To do this, we fit our sstGPLVM accounting for sample sex from Yoruba iPSCs (8, 24) and also a model with no fixed covariates. The high-dimensional gene counts are fully separable into sex by logistic regression (area under the curve (AUC) = 1.0), due to the presence of X- and Y-linked genes that will be differentially expressed across sexes. The addition of a fixed sex covariate in the low-dimensional space reduces the separability from an AUC of to 0.43, which shows a reduction in the confounding effect of sex on transcription after correction. Visually, we can see greater mixing of the male- and female-derived cells with the inclusion of a fixed sex covariate (Figure 3).

**Fig. 3.**
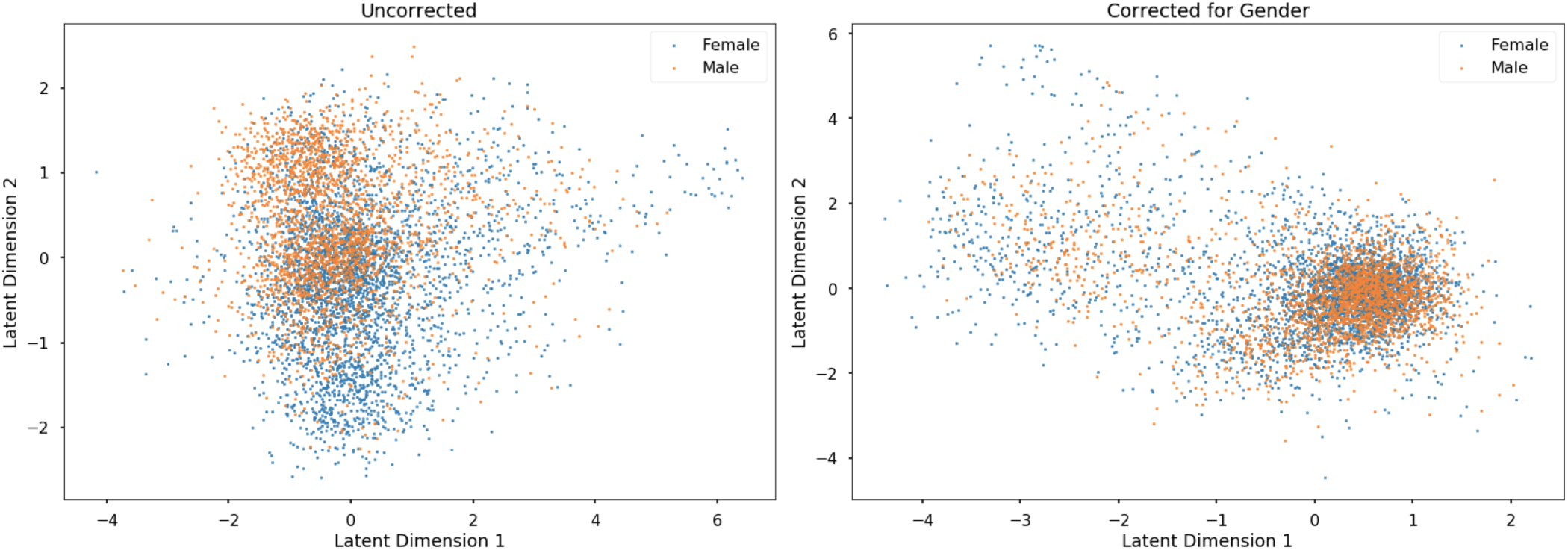
Reduction of sex separation in Yoruban stem cells. First two latent dimensions of sstGPLVM embedding of Yoruban stem cells (left) without correction, and (right) correcting for sex.

We can use the latent space to project back to high dimensional space as if all the cells came from the same sex. When we impute counts as if all cells were male-derived, we see an increase in expression of the switched cells in Y-linked genes *RPS4Y1*, while average expression of X-linked genes such as *USP9X* decreases (Additional File 2). Alternatively, when we impute counts as if all cells were female-derived, we see an increase of X-linked genes such as *BEX3, TMSB4X*, and *PRDX4* in switched cells.

### Aligning scRNA-seq with spatial seqFISH_+_ data

Next, we used sstGPLVM to jointly model scRNA-seq expression data with seqFISH_+_ expression and spatial information (25, 26). Data from seqFISH_+_ consists of cell specific counts of 10,000 genes and cell centroid coordinates for each cell. This scRNA-seq data contains counts for 24,057 genes. We used the (*x, y*) coordinates of the cell centroids from the seq-FISH+ processed results as fixed variables for seqFISH_+_ expression samples but missing for the disassociated cells assayed using scRNA-seq; we also added two latent dimensions for all cells. We incorporated a binary variable indicating seqFISH_+_ or scRNA to account for different levels of expression in each modality. We fit scRNA-seq data from mouse brain (26) with spatial information from the mouse olfactory bulb, cortex, and subventricular zone (SVZ) using 9,541 shared genes (25).

Our model is able to infer the spatial coordinates of each of the scRNA-seq cells with respect to the three seqFISH_+_ brain regions. We explored the organization of the scRNA-seq cells with respect to their inferred positions (Figure 4). We used mutual nearest neighbors to identify inferred “adjacency” of disassociated cells from scRNA-seq. With given cell type labels, we compared the frequency of neighboring types relative to distribution of cell types in the sample. A Fisher’s exact test of the “adjacency” table indicates that the organization is unlikely to occur by chance (*p* ≤ 0.0005). In the olfactory bulb, we observed that oligodendroctyes are 1.8 times as likely to be adjacent to astrocytes relative to astrocyte’s base frequency; conversely, astrocytes are 1.8 times as likely to be near oligodendroctyes relative to their own frequency (Figure 5). A potential reason is that astrocytes are responsible for promoting myelination activity of oligodendrocytes (27). The original seqFISH_+_ data showed a similar relative contact frequency between cluster 10 astrocytes and cluster 25 oligodendrocytes. Similarly, microglia are 1.4 times as likely to be adjacent to endothelial cells relative to the base frequency of microglia. This enrichment was also observed in the original analysis of seqFISH_+_ olfactory bulb organization (25).

**Fig. 4.**
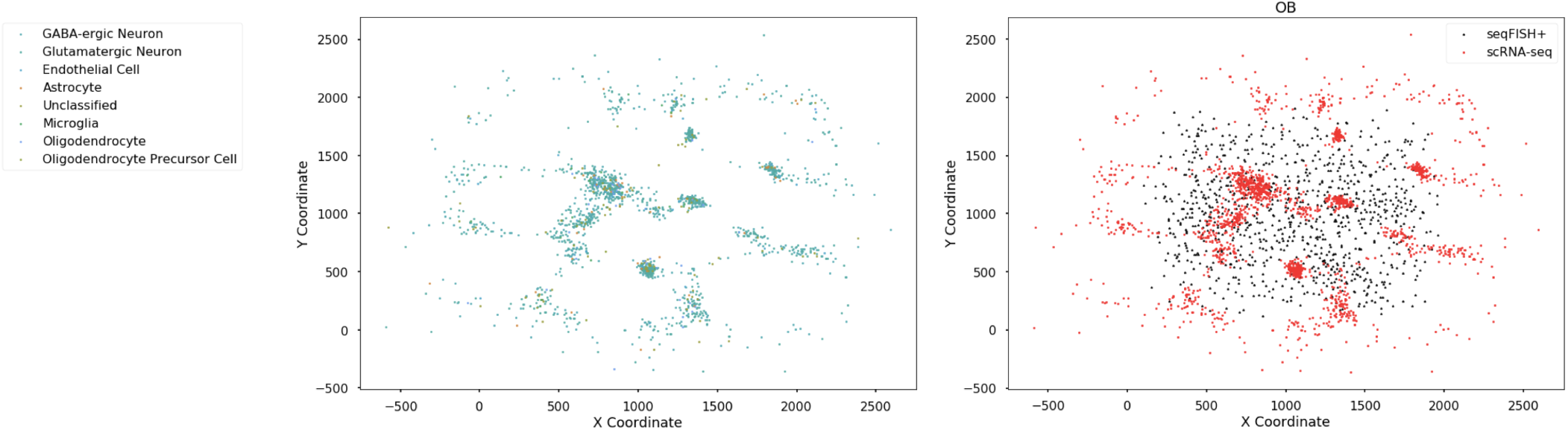
Joint spatial embedding of mouse brain cells. (Left) Inferred spatial coordinates of scRNA-seq data colored by cell type. (Right) Spatial coordinates from seqFISH_+_ (black) and estimated spatial coordinates for scRNA-seq data (red).

**Fig. 5.**
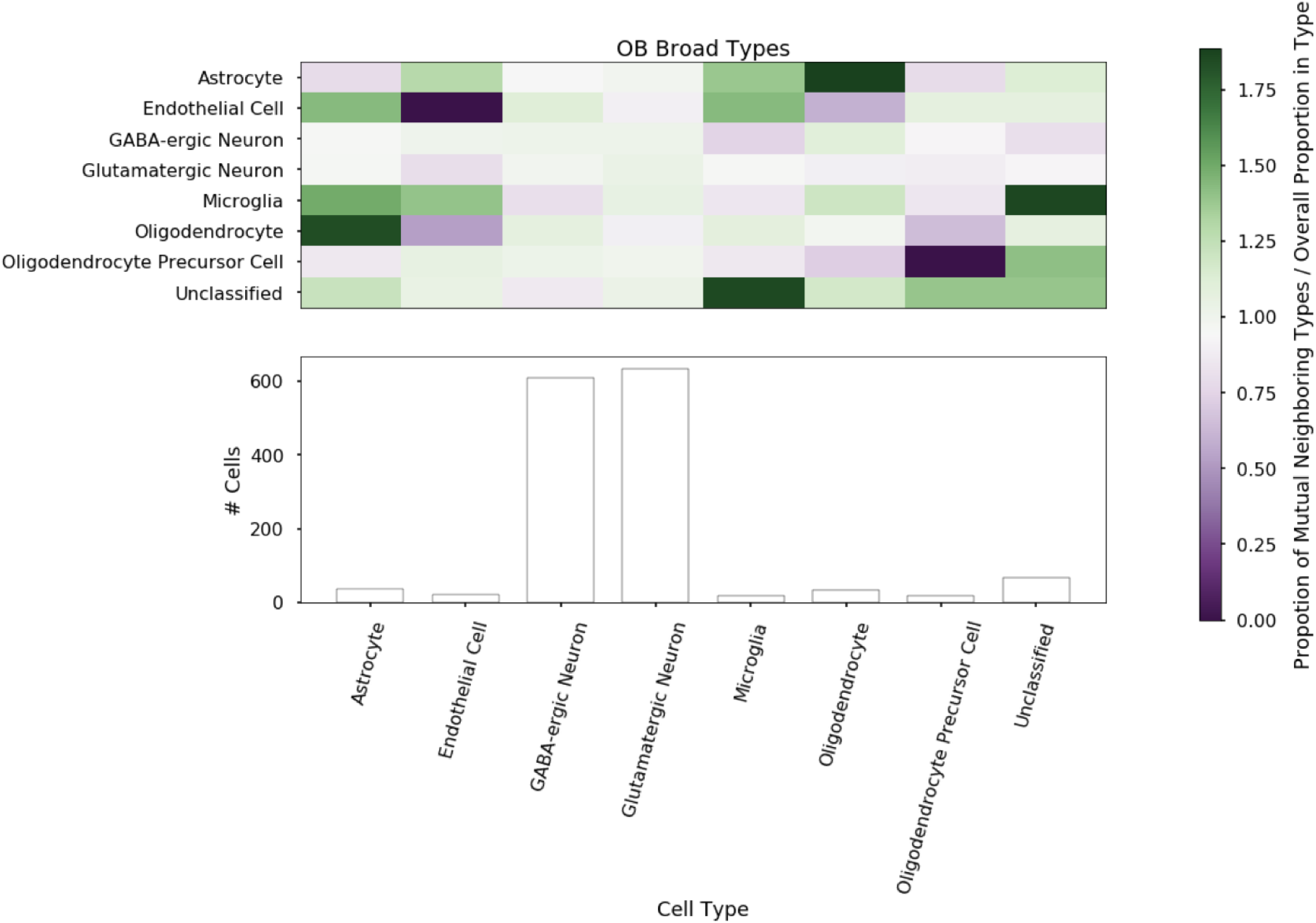
Spatial organization of the olfactory bulb broad cell types. (top) Ratio of percent nearest neighbors of each broad cell type relative to abundance in the olfactory bulb. Rows indicate the cell’s own types and columns represent the neighbors’ types (bottom). Abundance of cell types aligned inside seqFISH_+_ coordinates.

In our inferred location data, we examined enrichment among the higher-resolution cell labels. We observed that L2/3 Ptgs2 cells are enriched for adjacency to L4 Scnn1a cells (Figure 6). However, we note that these counts tend to be smaller and thus the results are less reliable. In the SVZ, we observed that endothelial cells and microglia are more likely to be self adjacent rather than adjacent across types (Figure 7). The cortex shows patterns similar to the SVZ, but with less self-adjacency (Figure 8). We noticed that GABA-ergic neurons and Glutamatergic neurons, the primary two cell types in these mouse brain regions, tend to be more self-adjacent in the cortex and SVZ than in the olfactory bulb in both the inferred spatial single cell data and the original seqFISH_+_ data. Given inferred cell adjacency, we wanted to quantify how spatial organization affects gene expression. To do this, we compared gene expression within cell types across different cell-cell interactions. The full scRNA-seq data allowed us to identify more genes that are spatially differentially expressed than the seqFISH_+_ counts. Astrocytes that are adjacent to oligodendrocytes in the olfatory bulb, for examples, express more *Espnl* that astrocyes that are not adjacent to oligodendrocytes (*p* ≤ 0.004). *Espnl*, a paralog of the gene that encodes the ESPIN protein, is associated with actin bundling and the neural process of hearing (28, 29). We observe a clear decay in expression of *Epsnl* when comparing expression levels to the distance of the cell to the nearest oligodendrocyte (Spearman’s *ρ* = −0.37, *p* ≤ 0.036; Figure 9). In oligodendrocytes that border astrocytes, we also found increased expression of genes involved in neural pathways such as *Cckar* (*p* ≤ 0.003) (30) and *Manf* (*p* ≤ 0.003) (31). We found a drop in *Cckar* expression in oligodendrocytes as a function of the distance to an astrocyte (Spearman’s *ρ* = −0.50, *p* ≤ 0.002; Figure 9). Moreover, using a normal approximation to the t-distribution, we can estimate the counts of the genes that are missing from the seqFISH_+_ data but present in the scRNA-seq data (e.g., *SNAP25, VSNL1*; Additional File 3)

**Fig. 6.**
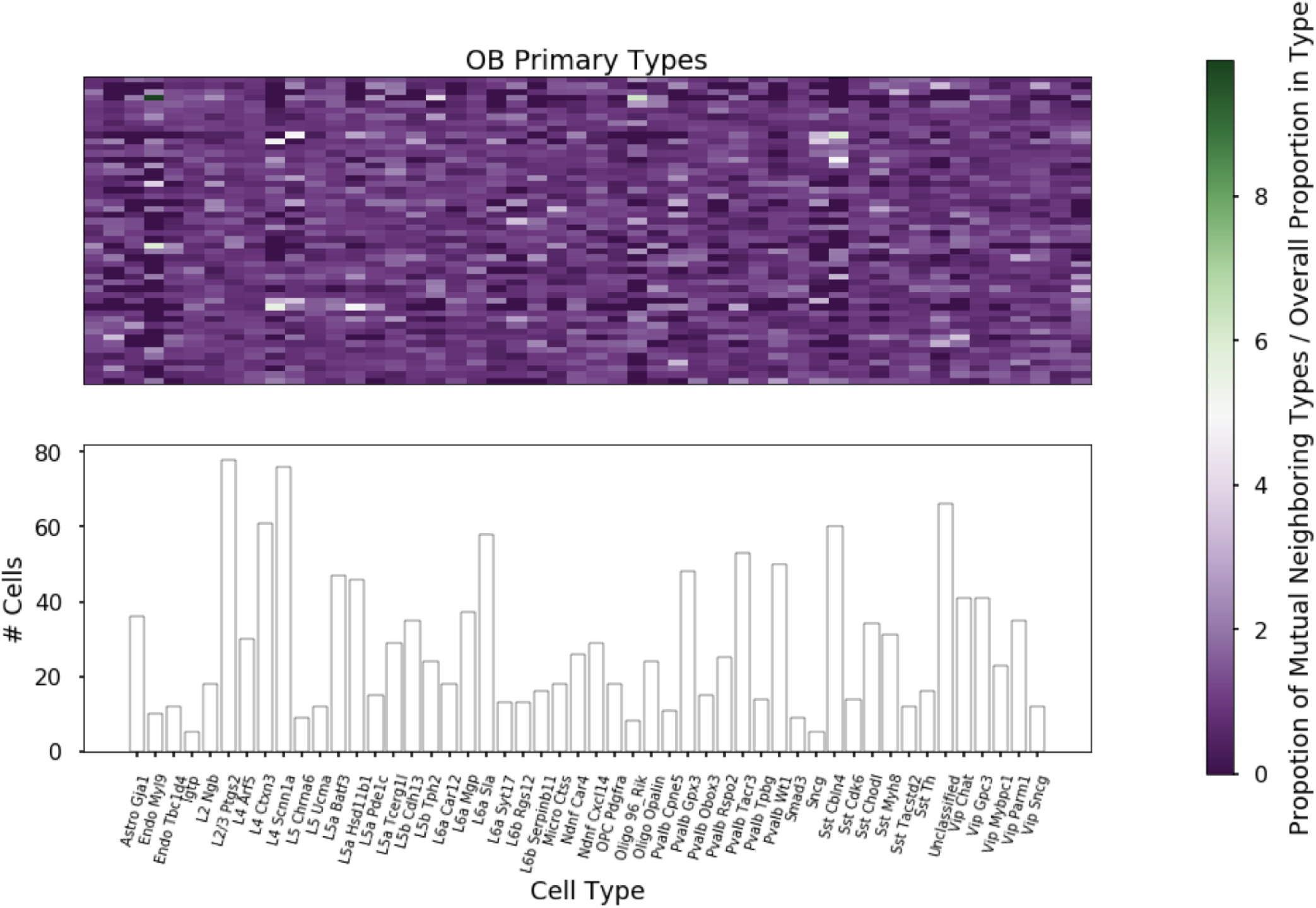
Spatial organization of the olfactory bulb primary cell types. (top) Ratio of percent nearest neighbors of each primary cell type relative to abundance in the olfactory bulb. Rows indicate the cell’s own types and columns represent the neighbors’ types (bottom). Abundance of cell types aligned inside seqFISH_+_ coordinates.

**Fig. 7.**
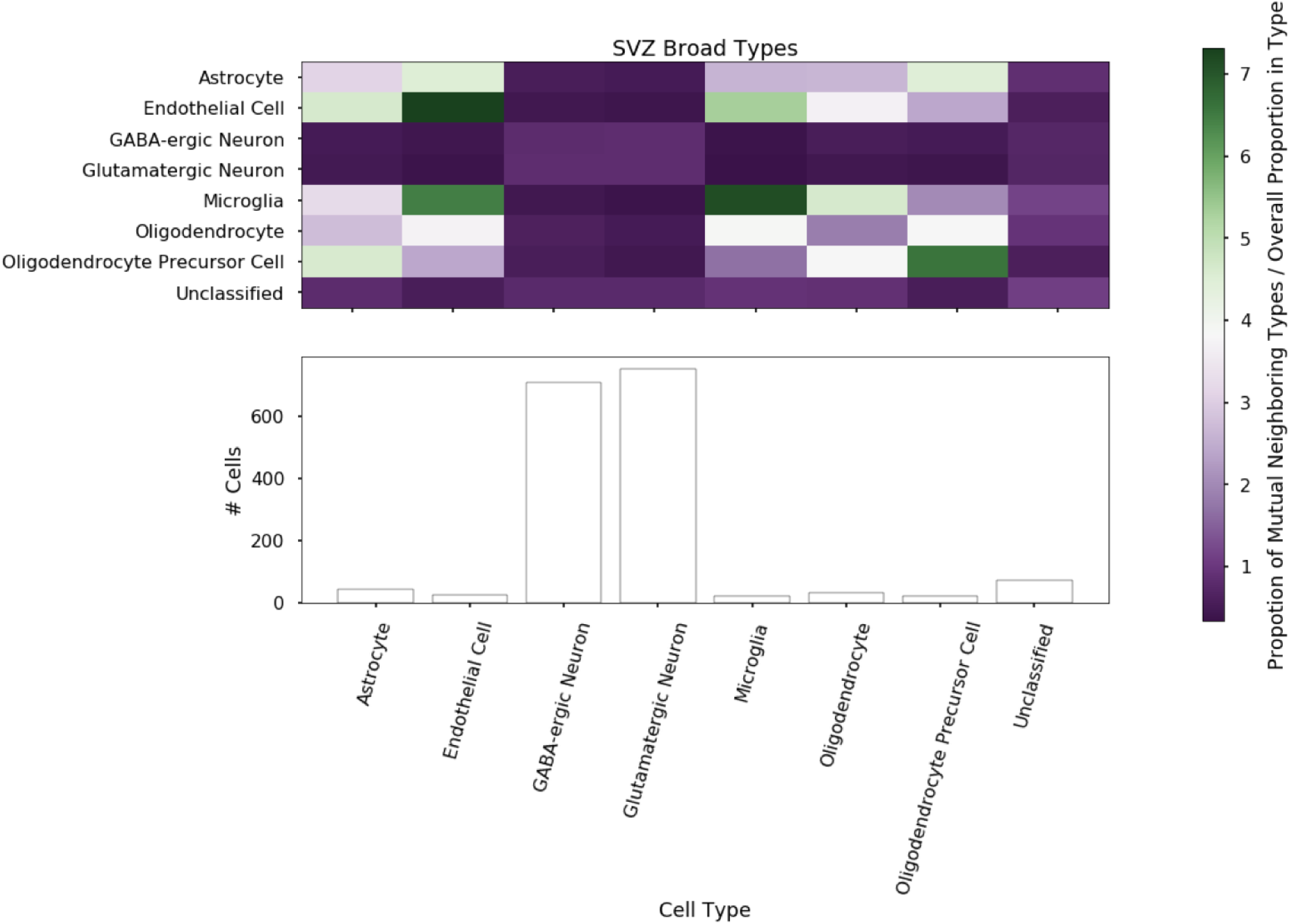
Spatial organization of the SVZ broad cell types. (top) Ratio of percent nearest neighbors of each broad cell type relative to abundance in the subventricular zone. Rows indicate the cell’s own types and columns represent the neighbors’ types (bottom). Abundance of cell types aligned inside seqFISH_+_ coordinates.

**Fig. 8.**
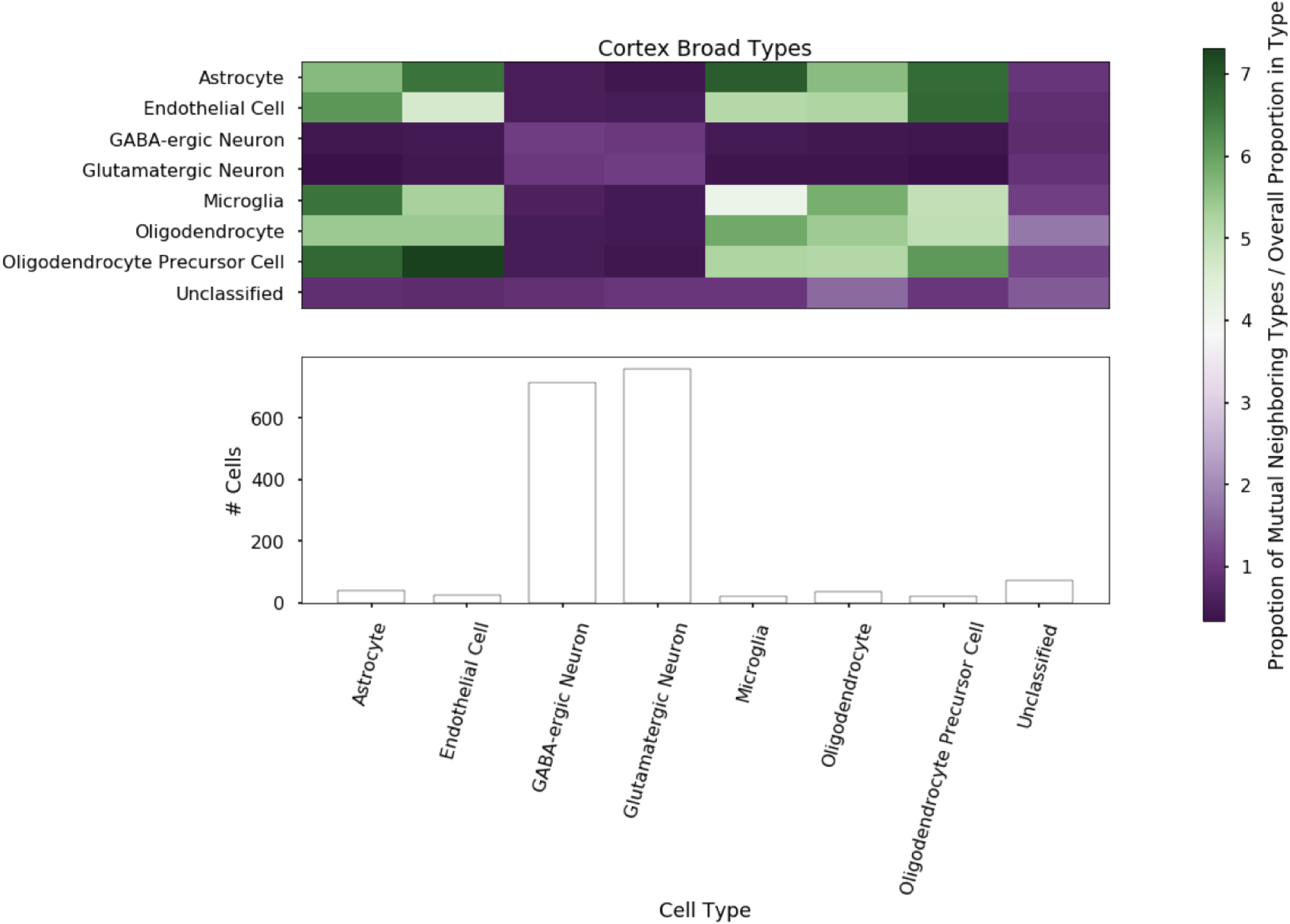
Spatial organization of the cortex broad cell types. (top) Ratio of percent nearest neighbors of each broad cell type relative to abundance in the cortex. Rows indicate the cell’s own types and columns represent the neighbors’ types (bottom). Abundance of cell types aligned inside seqFISH_+_ coordinates.

**Fig. 9.**
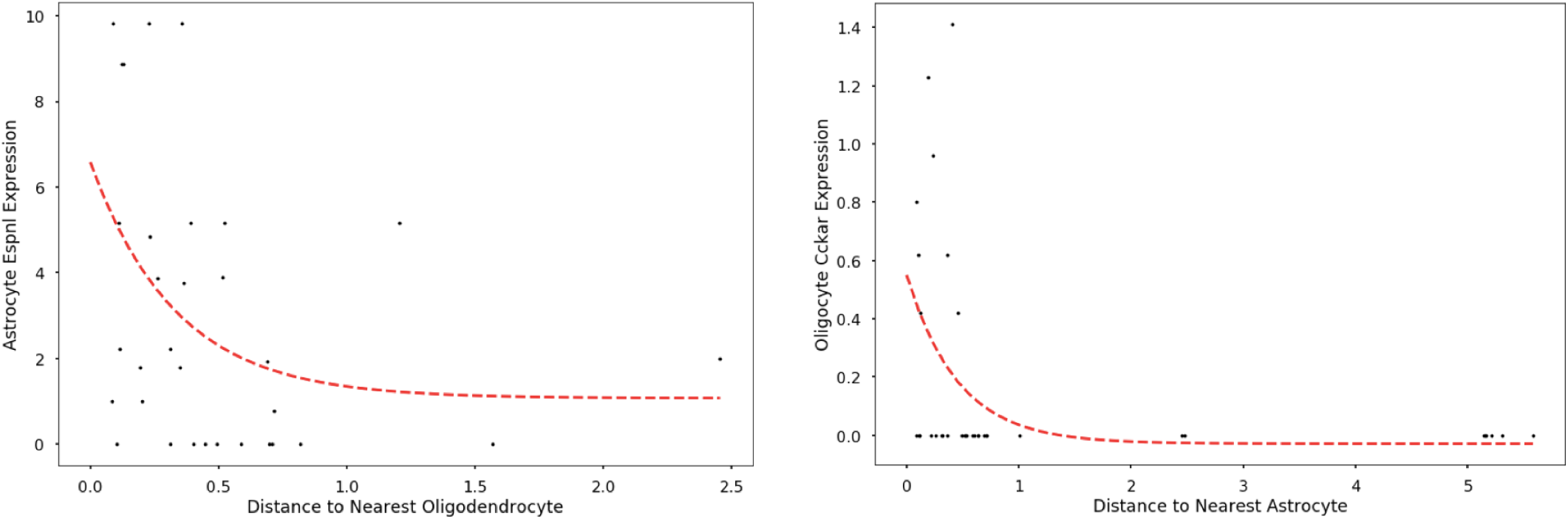
OB Gene expression across space. (left) Expression of *Espnl* in scRNA-seq astrocyte cells as a function of distance to nearest oligodendrocyte (bottom) Expression of *Cckar* in scRNA-seq oligodendrocyte cells as a function of distance to nearest astrocyte

The ability of sstGPLVM to fill in missing or unobserved co-variates such as spatial coordinates allows the integration of multimodal data for richer insights into single cell characteristics. New technologies often capture information on fewer genes or cells at time-of-development. The ability to augment data from these spatial technologies with richer count data from scRNA-seq with disassociated cells improves the power for discovery within these new techniques.

## Discussion

We developed a flexible alignment method for single cell RNA-seq data. Our method, sstGPLVM, has four types of behaviors that enable robust and flexible analysis of single cell data sets. First, sstGPLVM provides a mapping between high and low dimensional spaces that removes variation due to known covariates such as batch or sex. In our results, we showed the impact of this with respect to clustering and also imputation as compared to existing state-of-the-art methods. Second, sstGPLVM provides uncertainty estimates in the alignment of nonlinear manifolds. We demonstrate that these uncertainty estimates provide meaningful information about the accuracy of the embedding of each cell and can be propagated effective to downstream analyses. Third, sstGPLVM provides reference-free regularization that preserves variation from sources other than the fixed covariates. While there is no ground truth “counterfactual” counts for a cell sequenced under different conditions, we can project the latent space to gene space using a particular value of the fixed co-ordinates to approximate the counts if all cells were from the same experiment. Finally, sstGPLVM allows the underlying cell type proportions to vary substantially across batches. In both the simulated and pancreas cell data, cell types were not present in all batches, yet the model was able to effectively learn the latent structure. sstGPLVM is able to integrate single cell information across biological and technical conditions, and is flexible to different types of covariates. While current methods such as MNN and CCA account solely for the discrete batch variable, sstGPLVM can be used with continuous information such as spatial location or biological covariates such as sex. With respect to spatial locations as a fixed covariate, while our results here show promise using a single image (with a single coordinate system) for each projection, the relative (*x, y*) coordinates were local to that image; in the future, development of a common coordinate framework (CCF) (32) for describing the relative locations across images would be useful in integrating data using shared spatial landmarks. As new methods provide richer data about cell phenotypes in conjunction with gene expression, such as methylation or protein signals, sstGPLVM will be a powerful tool to work across modes and with multiple data representations.

## Conclusions

We demonstrate the ability of semi-supervised tGPLVMs to integrate multiple single cell data sets for joint analysis. sstGPLVM is better at recovering true manifolds than existing methods such as MNN and CCA as demonstrated on simulated data with ground truth. We see that uncertainty estimates of embeddings are useful when jointly analyzing pancreas cells to identify which cell’s embeddings can confidently be used for downstream analyses. Finally, we demonstrate that sstGPLVM can be used across modalities by jointly fitting spatial seqFISH_+_ and scRNA-seq to learn spatial mappings for dissociated single cells. With this knowledge, we can uncover genetic patterns and patterns across space that may not be accessible in one of the modalities separately. As single cell technologies evolve, the ability to combine data sets will become more vital part of any analysis pipeline. The sstGPLVM offers a principled method for integrating single cell data and potentially other types of multi-modal data.

## Notes

#### Summary of Updates

Updated methods section for clarity

https://github.com/architverma1/sc-manifold-alignment

