## Additional File 1 for "A Bayesian nonparametric semi-supervised model for integration of multiple single-cell experiments"

**Additional File 1: Pancreas Hyperparameter Optimization**

**Table 1: Hyperparameter Optimization for sstGPVLM Pancreas fit.** Using the sum of length scales for the Matérn ½ and RBF kernels, dimensions were ordered from most (lowest sum) to le ast (highest sum)"important." One repeat of K-means clustering was performed with K between 5 and 10, and evaluated using Adjusted Rand Score (ARS) and Normalized Mutual Information ( NMI)

| **Dimensions Included** | **ARS** | **NMI** |
|---|---|---|
| 1 | 0.12 | 0.22 |
| 2 | 0.14 | 0.24 |
| 3 | 0.27 | 0.36 |
| 4 | 0.31 | 0.39 |
| 5 | 0.34 | 0.42 |
| 6 | 0.32 | 0.41 |
| 7 | 0.31 | 0.41 |
| 8 | 0.32 | 0.42 |
| 9 | 0.24 | 0.33 |
| 10 | 0.19 | 0.29 |
