## Additional File 2 for "A Bayesian nonparametric semi-supervised model for integration of multiple single-cell experiments"

**Additional File 2: Additional Sex Imputation Results**

**Supplemental Table 1:** Genes that show the most change in expression when imputed to opposite gender.

| **Male to Female** | | **Female to Male** | |
|---|---|---|---|
| **Increase** | **Decrease** | **Increase** | **Decrease** |
| BEX3 | NUP98 | RPS3 | NUP98 |
| ACTB | ATP6 | GAPDH | H4C3 |
| RPS4X | RPS4Y1 | RPS18 | BEX3 |
| TCEAL8 | H4C3 | RPLP1 | VMP1 |
| FTL | RPS19 | RPS4Y1 | CYTB |
| SMS | MT1X | RPS6 | ACTB |
| PRDX4 | IFITM1 | RPS25 | COX2 |
| TCEAL9 | ND4 | RPL31 | FTL |
| TMSB4X | RPS3 | RPL13A | ND4 |

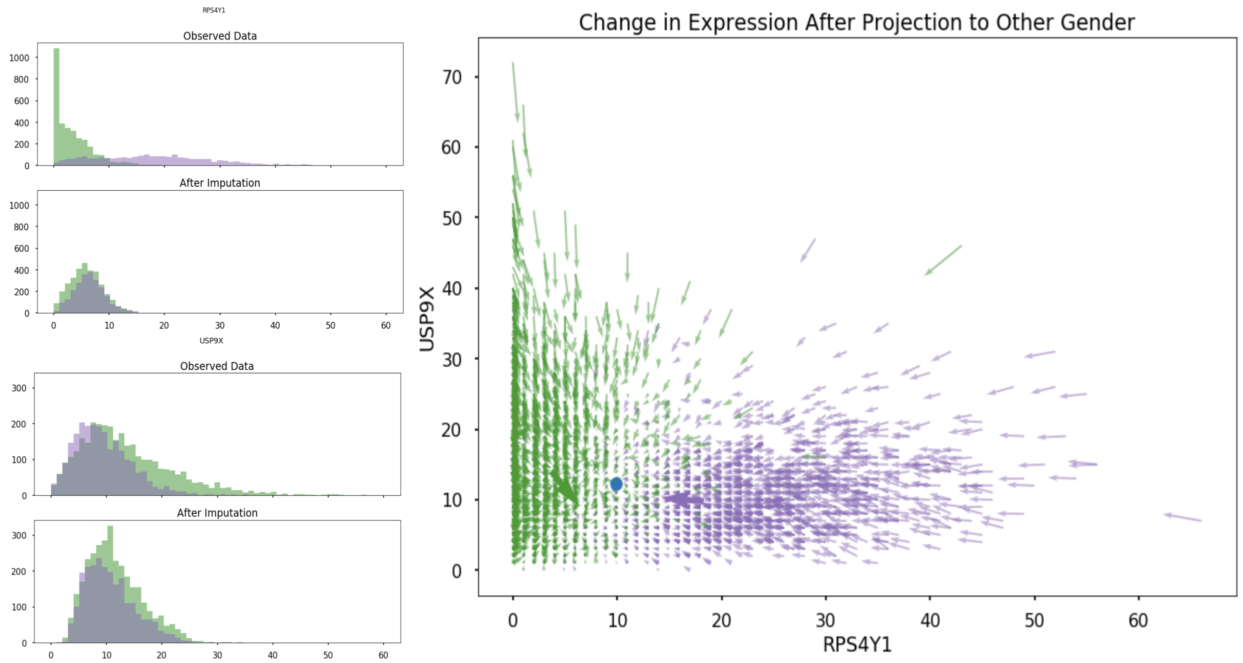


**Supplementary Figure 1:** (left) Distribution of RPS4Y1 and USP9X in male (purple) and female (green) cells in original data and after imputation to opposite sex (right) Quiver plot where thin arrows indicate the change from original observation to imputed to opposite sex, thick arrows represent the mean change, and blue represents the sample mean.
