## Additional File 3 for "A Bayesian nonparametric semi-supervised model for integration of multiple single-cell experiments"

### Additional File 3: Imputation of genes not available in seqFISH+ data

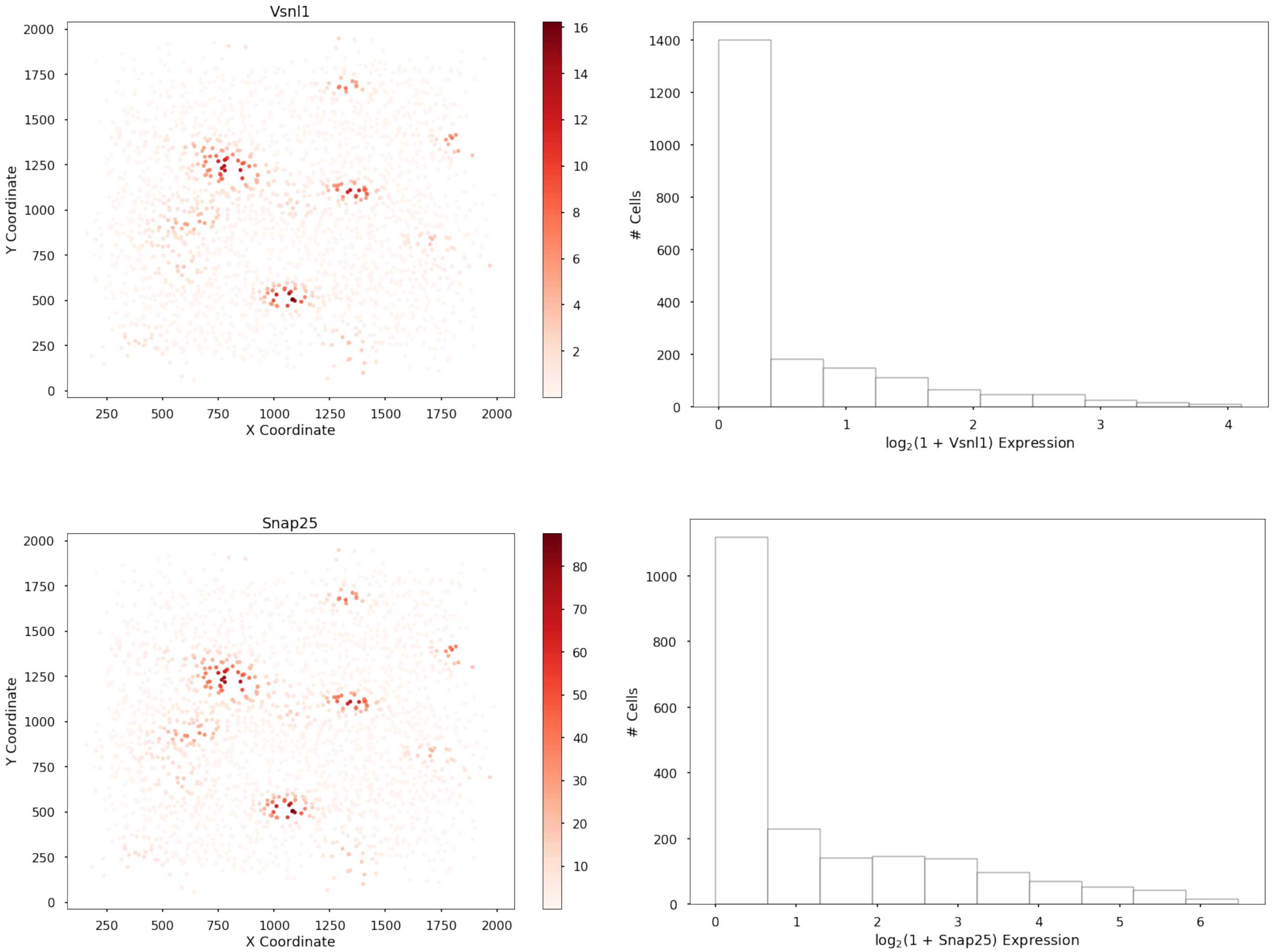


**Supplementary Figure 1:** Spatial visualization and histograms of imputed counts of *VSNL1* (top) and *SNAP25* (bottoms) genes not available in the original seqFISH+ data.
